# Fly, land, listen: Autonomous intermittent locomotion enables scalable low-noise drone ecoacoustic surveys

**DOI:** 10.64898/2026.09.28.754547

**Authors:** Milica Ostojic, Oscar Pang, Matt Phelps, Mirko Kovac, Sarab S. Sethi

**Affiliations:** Department of Life Sciences, Imperial College London, Imperial White City Campus, London, UK W12 0BZ; Department of Aeronautics, Imperial College London Imperial South Kensington Campus, London, UK SW7 2AZ; Knepp Estate Rewilding Project, The Estate Office, Knepp Castle West Grinstead, West Sussex, UK RH13 8LJ

**Keywords:** Ecoacoustic surveys, biodiversity monitoring, unmanned aerial vehicles, autonomous drones, drone noise, bird vocalisations, open-source robotics, sustainability robotics

## Abstract

1. Ecoacoustic monitoring is enabling scientists and land managers to monitor and manage biodiversity more effectively and cost-efficiently in the face of human pressures and rapidly changing climates. Currently, most ecoacoustic surveys use manually deployed static sensors to record data, limiting the scale and reach of surveying efforts. Here we present a proof-of-concept autonomous drone platform that can use intermittent locomotion to conduct ecoacoustic surveys using an onboard sensor.
2. Our custom prototype is able to fly, navigate, and avoid obstacles autonomously, land at a pre-determined location, record audio from an onboard microphone whilst static, before taking off and moving to the next sampling site. Autonomous navigation and operation enable greater sampling flexibility, reach, and scalability. Furthermore, by recording audio only whilst landed, noise from the drone’s rotors does not mask signals or disturb animals, simplifying signal processing and downstream ecological analyses.
3. We conducted trials in a scrubland habitat at the Knepp Estate in West Sussex, where our prototype demonstrated successful autonomous navigation and obstacle avoidance. Furthermore, we found that avian biodiversity data collected from the drone platform was comparable to that from traditional static acoustic sensor deployments, and that vocalisation patterns were not significantly impacted by the noise of the drone arriving or leaving a site.
4. While scaled deployments of our technology would require further technical and regulatory challenges to be solved, our first demonstration of autonomous intermittent robotics-assisted ecoacoustic surveys lays the foundations for more cost-effective and far-reaching biodiversity surveys, with transformative potential for conservation, agricultural management, biosecurity, and more.

## 1. Introduction

Reliable biodiversity monitoring is becoming ever more imperative as pressure on global ecosystems increases. While monitoring efforts are slowly scaling up, there are still significant spatial (1) and temporal (2) gaps in biodiversity data globally. Filling these data gaps is essential to drive effective policy and conservation work, as is monitoring the impact of interventions, such as policy changes (3) or rewilding efforts (4) to evaluate outcomes and inform future designs. To achieve this wide-spread, consistent monitoring, a change in approach is needed (5). Traditional monitoring techniques, such as tagging (6), manual point counts and transects (7), can be considered invasive, as well as labour and resource intensive (7). Alternative, automated methods are therefore gaining in popularity due to their ability to increase the scale, cost-effectiveness and time-efficiency of monitoring, allowing scientists and land managers to monitor environmental health across greater spatial and temporal scales with improved accuracy.

A promising approach to rapidly surveying biodiversity is to use acoustic data (hereafter, ecoacoustics) (8). When large-scale audio recording and automated data analysis techniques are combined, as is the case with passive acoustic monitoring (PAM), ecoacoustics can be used to obtain invaluable information about vocal species and their environments (7). Compared to traditional biodiversity surveys, PAM is less invasive, less labour intensive and can allow species to be surveyed at very large spatiotemporal scales and fine resolutions (9–11). Nevertheless, a significant amount of manual labour is still required to deploy, maintain (e.g., replacing batteries and retrieving SD cards), and retrieve sensors during PAM surveys (10). Therefore, deploying large sensor networks or surveying remote inaccessible regions can be logistically complex, costly, and even dangerous. Storage limitations can be mitigated by continuously transmitting data over a mobile data network (12) or long-distance radio (13), and solar panels can resolve the need for battery replacement (13). However, even when using off-grid powered and networked sensors, initial deployment and final retrieval are still unavoidable.

Using robotic platforms, particularly Unmanned Aerial Vehicles (UAVs) (commonly known as drones) (14,15), to collect PAM data offers a way to sidestep the manual labour required for sensor deployment, maintenance, and retrieval. Existing ecoacoustic drone studies have used drones to carry microphones hanging from a rope over short periods of time (i.e. minutes) (16), with relative success for detecting bird vocalisations, although drone noise meant certain species were underrepresented (17). Hardware engineering efforts to reduce drone motor noise exist (18), but it seems unlikely that it will ever be eliminated entirely.

Drone noise can alternatively be attenuated when post-processing audio (19). However, continuous and extended periods of noise do not only affect audio recordings, but also the wildlife itself as species can demonstrate avoidant behaviour (16). Furthermore, recording from drones in flight is significantly limited by battery life, meaning that only short snapshots of data can be collected (17). Finally, existing drone-based ecoacoustic studies have used both manual (20) and semi-autonomous operation (21,22). Fully autonomous operation with obstacle avoidance that integrates acoustic biodiversity data has not been demonstrated yet, limiting the accessibility and autonomy of using drones in PAM surveys.

A potential drone-based ecoacoustic survey design that can mitigate the limitations of PAM is intermittent locomotion (23,24), which would involve the drone flying between sampling locations and only recording while landed at each of them. Drones that use this strategy to “hop” between sites offer many potential advantages over existing approaches. Audio can be recorded for extended periods while a drone is landed at a site, providing more than just short snapshots of biodiversity data. The spatial resolution and reach of surveys would be increased, as individual sensors don’t need to be manually deployed and retrieved, and areas which may be difficult to access on foot can be accessed by air. Real-time data could even make adaptive sampling (25) feasible, as moving during a deployment becomes straightforward. Furthermore, maintenance burden is reduced as only one drone is serviced rather than several static recording devices. Not having simultaneous recordings from multiple sites (as a network of static recorders would deliver) may result in intermittent locomotion being less effective for surveys of rare species (26), but simulations have shown that intermittent locomotion can still deliver reliable biodiversity data for many species and ecological questions (26).

Here we introduce novel designs for an autonomous drone with obstacle avoidance, which uses intermittent locomotion to collect PAM data and survey biodiversity. Our modular design uses open-source components and custom 3D printed parts to present a more purpose-built and flexible (27) alternative to off-the-shelf drone models. Alongside our technical designs, we present the results of field trials at the Knepp Estate in West Sussex, UK (28), where we demonstrate how our drone can be used for autonomous avian biodiversity surveys.

## 2. Hardware design

### 2.1 Hardware overview

Our drone uses a Pixhawk 6C flight controller, a quadcopter airframe of diagonal length 450mm with four 2216-920KV Brushless Motors, 20A 3-4S Electronic Speed Controllers (ESCs), and four self-locking propellers. The main source of power is a 3000mAh 4S LiPo Battery, which connects to the ESCs, the flight controller and an onboard companion computer via a Power Distribution Board (PDB).

The flight controller runs the open-source PX4 firmware which commands the motors (via the ESCs), onboard sensors, and the companion computer. The sensors connected to the flight controller include a GPS module that provides localisation data, a built-in IMU that measures the linear and angular acceleration of the drone, a radio telemetry module that receives and sends live telemetry data to the ground control station (which can be a laptop or phone running the QGroundControl application), a radio receiver that gets drone pilot commands from a radio controller fitted with a radio transmitter and long-range antenna, and an altimeter (we use a separate range finder as opposed to the built-in flight controller barometer) that collects altitude data. The flight controller also translates control commands from the pilot or an autonomous algorithm to the required speed of the motors by sending signals to the ESCs via a pulse-width modulation (PWM) module.

The NVIDIA Jetson Orin Nano onboard computer, which is running autonomous obstacle avoidance software, is connected physically to the flight controller via an adapter that converts from USB to the TTL UART interface (Transistor-transistor logic, Universal Asynchronous Receiver-Transmitter) and is powered by the LiPo battery via the PDB. The computer is flashed with the Jetpack 5.1.2 OS, which is based on Ubuntu 20.04. An Intel RealSense D455 stereo depth camera is connected to the Jetson via USB to provide vision data (through depth mapping). MAVROS, a ROS (Robot Operating System) package, uses the MAVLink protocol to communicate between the obstacle avoidance software (which is built using ROS), the flight controller and the ground station. This includes communicating telemetry data (such as GPS status, attitude, battery level etc.) and commands (both manual commands from a drone pilot and autonomous software path planning).

An AudioMoth mounted above the Jetson captures acoustic data. The full assembly is presented in Figure 1, with a list of components and costs in Table 1 (models used and links to purchase can be found in the Supplementary Table), a network of components shown in Figure 2, and further details on key components in the following subsections.

**Figure 1:**
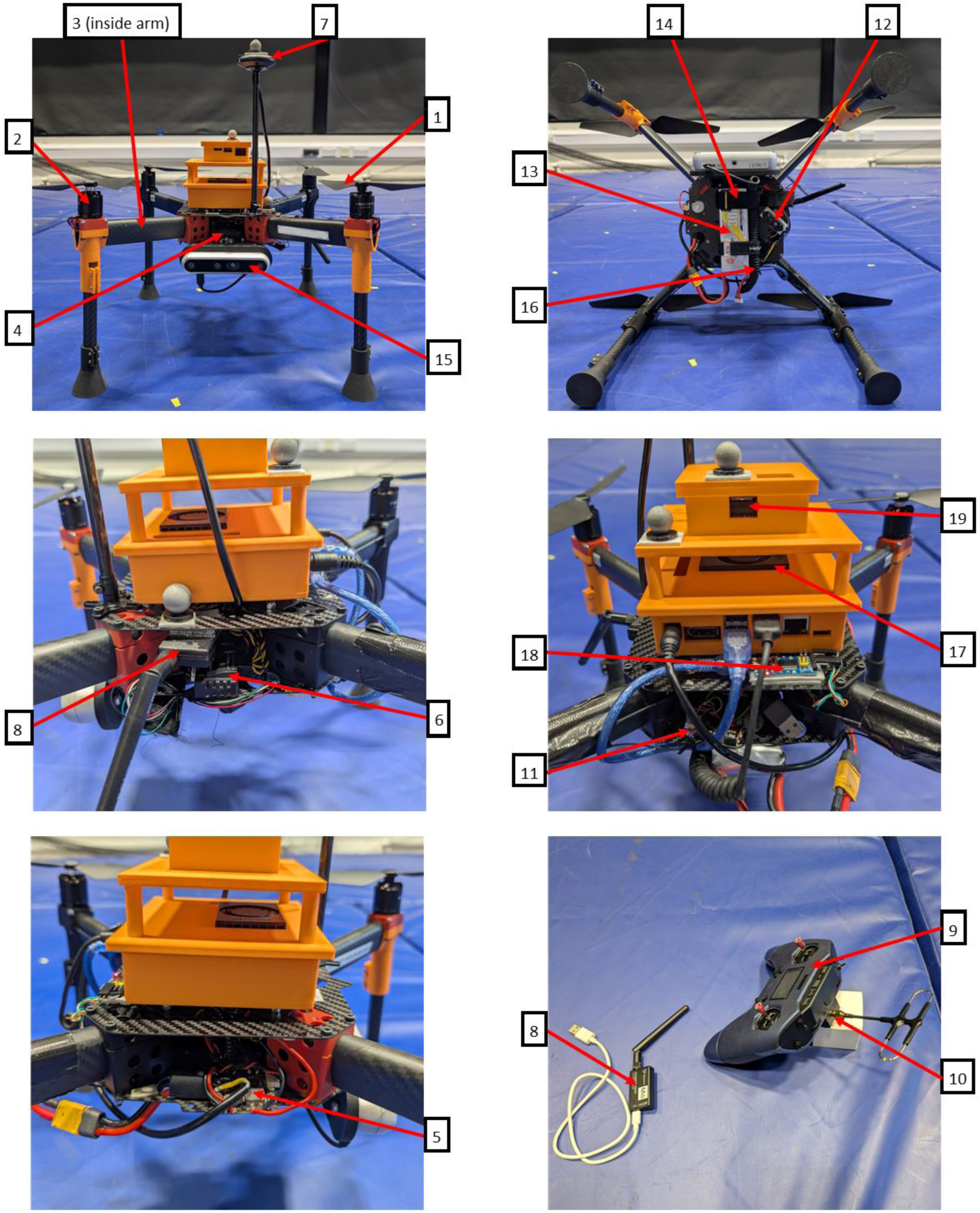
Images of multiple orientations of the drone with labels corresponding to the components, as detailed in Table 1.

**Figure 2:**
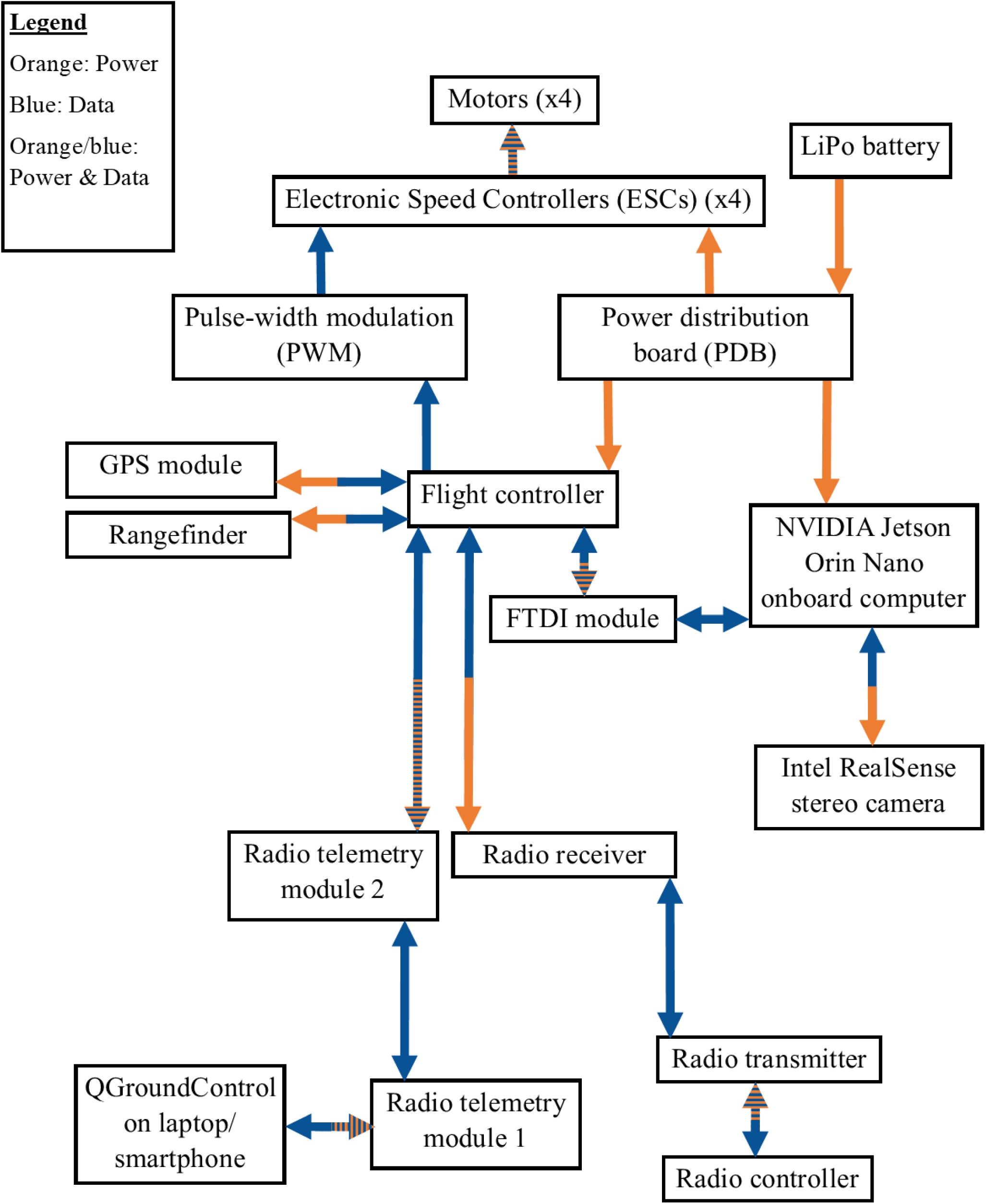
Network of components, showing the flow of power (orange) and data (blue).

**Table 1:** List of components involved in the drone design and their costs, corresponding to the labels in Figure 1.

| Label | Component | Cost |
| --- | --- | --- |
| **N/A** | Drone frame | £239.00 |
| **1** | Propellers (x4) | £7.99 each |
| **2** | Brushless motors (x4) and bullet connectors | £19.90 each |
| **3** | Electronic Speed Controllers (ESCs) (x4) | £19.99 each |
| **4** | Flight controller | \$186.98<br>(£137.04)<br>Price for items 4-6 included |
| **5** | Power distribution board (PDB) | \$20.99<br>(£15.39)<br>When purchased separately |
| **6** | Pulse-width modulation (PWM) board | \$26.99<br>(£19.78)<br>When purchased separately |
| **7** | GPS module | \$54.99<br>(£40.31) |
| **8** | Radio telemetry | \$58.99<br>(£43.65) |
| **9** | Radio controller | £128.26 |
| **10** | Radio transmitter | £69.90 |
| **11** | Radio receiver | £32.90 |
| **12** | Rangefinder | £44.00 |
| **13** | LiPo Batteries | £33.43 each |
| **14** | Additional battery strap | £2.50 |
| **15** | Stereo camera | \$419.00 (£310.18) |
| **16** | Shielded USB cable for stereo camera | £11.99 |
| **17** | Onboard computer | £457.58 |
| **N/A** | Nvme internal storage device for onboard computer | (Discontinued) £63.78 /<br>(Alternative) £129.98 |
| **18** | FTDI module | £6.50 |
| **19** | Acoustic device | £79.99 |

### 2.2 Custom designs for modularity and adaptation

We designed and 3D printed (in PLA and PLA-CF) a number of components to modify the HexSoon – EDU-450 frame for our use-case (Figure 3).

**Figure 3:**
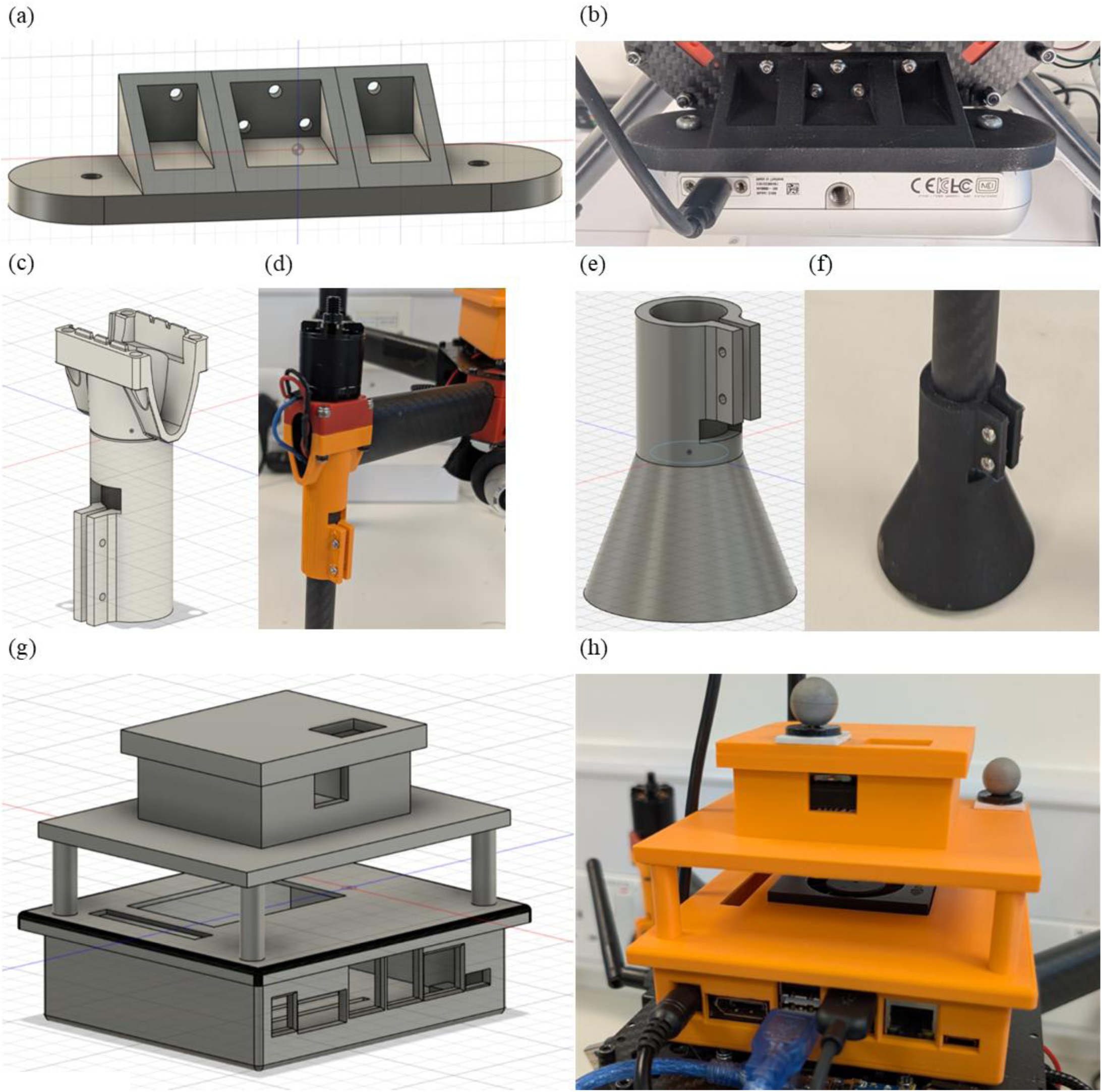
(a) CAD design for the Intel RealSense camera mount. (b) Intel RealSense camera mount printed and attached to the drone platform. (c) CAD design of the leg mount, adapted from a CAD file of the original drone kit frame. (d) Leg mount printed and attached to the drone platform. (e) CAD design of the leg base. (f) Leg base printed and attached to the drone platform. (h) CAD design of the Jetson Orin Nano and AudioMoth combined case. The Jetson case is adapted from a public design available on the Bambu Lab Online models repository. (g) Jetson Orin Nano and AudioMoth combined case printed and secured on the drone platform, with motion tracker markers attached.

#### 2.2.1 Intel RealSense camera mount

The Intel RealSense camera mount (Figure 3a) is designed to screw onto the front underside of the bottom plate through the existing holes in the plate. The camera can then be secured on the front of the mount through the two holes, matching the screw holes in the camera itself (Figure 3b). The camera is oriented in the same plane as the flight controller, providing depth measurements for the area in front of the drone as it flies. As the camera has a depth field of view (FOV) of 87° × 58° (29), this orientation enables obstacle avoidance for a wide area in front of the drone, preventing collisions in the direction of flight. Positioning the mount underneath the bottom plate also ensures that no parts of the drone, particularly the two frontal spinning propellers, are visible to the camera, which would otherwise interfere with the obstacle avoidance software. The camera mount was printed in PLA-CF for enhanced tensile strength.

#### 2.2.2 Landing gear

The original landing gear of the drone kit frame was unsuitable for our use-case, due to the very narrow radius of the leg poles leading to easy breakages and unstable landings on uneven ground. Therefore, we designed a new leg mount (Figure 3c), adapted from the original drone kit design, to accommodate a wider carbon fibre leg of radius 8 mm, with an even wider leg base (Figure 3e) on the other end to create a larger landing surface and improve the stability of the landing gear. The carbon fibre tube is secured in the leg mount and the leg base using screws through 2 mm holes (Figures 3d, f), used to ensure a tight seal and maximise stability. The leg mounts have been printed in PLA (Figure 3d), with the back legs in black and the front legs in orange to highlight flight direction from a distance. The leg bases were printed in PLA-CF (Figure 3f) for enhanced tensile strength. With our modular design, the leg mounts and leg bases can be adjusted or swapped out to create appropriate landing gear for different environments.

#### 2.2.3 NVIDIA Jetson Orin Nano and AudioMoth combined case

To secure the NVIDIA Jetson Orin Nano and AudioMoth on the drone, we designed a combined case (Figure 3g). The Jetson case was adapted from a public design available on the Bambu Lab Online models repository (30), with the addition of holes in the base that correspond to existing holes on the top plate of the drone frame, allowing the case to be attached to the top plate with screws, and minor adjustments to the size of the gaps in the case that correspond to the ports and the fan. The AudioMoth case was designed to be suspended by columns above the Jetson lid to allow for airflow over the Jetson fan. There are gaps in the sides of the AudioMoth case that correspond to the USB port, the SD card port, the switch, and the LEDs so that full functionality of the AudioMoth can be accessed without removing it from the case. The lid also has a gap over the microphone to reduce attenuation of the input audio. Unlike the standard setup of a static AudioMoth, the microphone is facing upwards. The lid of the AudioMoth case and the lid of the Jetson case are closed with snap fit joints, which ensure the Jetson and AudioMoth are securely sealed in their respective cases, even in crash scenarios. Motion tracker markers have been placed on the combined case (Figure 3h) to enable use of a VICON motion capture system for indoor position flight mode (where motion capture data is used for indoor positioning in the absence of outdoor GPS signal), for tests of stability and controllability of the drone. The combined case was printed in PLA, as its complex shape would not allow it to be printed in the more brittle PLA-CF. It was printed in orange to improve visibility of the drone from afar and increase chances of discovery and recovery should the case be detached from the drone in a crash.

In a trade-off between increased weight and direct draw on the drone battery, the AudioMoth mounted in the combined case is powered by its own batteries. This aspect of the design also allows the case to be easily adapted for other ecoacoustic recording devices or even other sensor modalities, that are similarly powered by their own batteries, by simply changing the dimensions and shape of the case suspended above the Jetson lid.

## 3. Drone software and operation

### 3.1 QGroundControl and Mission Planning

Operation of the drone in the field is achieved using QGroundControl connected to the radio telemetry. QGroundControl is an open-source software that presents all the live telemetry data about the drone, such as orientation and speed, flight mode, location, GPS service status and quality, flight controller internal logs, and battery level (31). It also allows the pilot to plan autonomous missions, enable obstacle avoidance and other fail-safe features, and calibrate the onboard sensors via the flight controller (31). It can be run on a laptop or a mobile device connected to a radio telemetry module, with the other module in the pair directly connected to the flight controller.

Outdoor autonomous missions are planned either directly through the QGroundControl interface (32) or by importing a mission file (33). The take-off and landing positions and waypoints in between must be provided, along with the height and position of each waypoint and the speed with which the drone should travel to each one. Once a mission has started, the drone will automatically switch to Mission mode and begin the mission following the set plan, using outdoor GPS position data for closed-loop control. If obstacle avoidance is enabled, the autonomous algorithm (see Section 3.2) will plan a trajectory around any obstacles to reach each waypoint.

### 3.2 Obstacle avoidance

In order to enable autonomous navigation outdoors with obstacle avoidance, we deployed the open-source PX4 Avoidance software (34) on the NVIDIA Jetson Orin Nano onboard computer, which uses the depth measurements from the stereo camera connected to the Jetson via USB to plan the path for the drone to take in order to avoid any obstacles.

The PX4 Avoidance project (https://github.com/PX4/PX4-Avoidance) uses depth mapping and the 3DVFH* algorithm to plan and update the trajectory around obstacles in outdoor environments (35). The 3DVFH* algorithm first builds a spherical histogram from current sensor information (36), with each histogram cell containing information on the distance to the closest obstacle in its area (36). The current histogram is then combined with a memory histogram that contains information from several previous timesteps (36), and this combined histogram is given to the A* algorithm (36), which generates nodes associated with each histogram cell and chooses a path forward based on which nodes are free and which path has the least cost (36).

The path consists of a series of waypoints, which are continually updated as new obstacles come into view.

These waypoints are provided to the Pixhawk 6C flight controller, which adjusts the orientation and the speed of the drone by changing the speed of the motors to capture the trajectory to the next waypoint. The flight controller also collates the current GPS position, the altitude from the rangefinder, and the acceleration and orientation from the IMU using its Extended Kalman Filter and sends the fused sensor data to the Jetson to update the obstacle avoidance software so it can change the path required to reach the set destination.

## 4. Operation and safety

### 4.1 Safe operation

We strongly recommend flying the drone manually outdoors in Position flight mode before attempting autonomous flight to ensure that the flight controller has been calibrated correctly and that the GPS signal is strong enough, which may take some time to achieve after powering on. Position mode can only be toggled if there is enough GPS signal, so if it is not possible, then autonomous operation will not be possible either. GPS signal received by the drone antenna can be affected by many factors, including position of the antenna, electromagnetic interference, physical obstructions in the surrounding environment (e.g. tree canopy, buildings, mountains), and availability of satellites. We also recommend attempting a mission without obstacle avoidance enabled first as a baseline test, in an outdoor area free of obstacles, to ensure that

Mission mode is working correctly – the drone should successfully follow the uploaded plan and should not significantly deviate from it.

During an autonomous flight, the drone pilot should stay within range of the drone to ensure that manual takeover with the radio controller is possible should the drone stray significantly from the mission plan, thereby reducing the risk of flyaway. When using PX4 firmware, by default, any stick movement on the radio controller during an autonomous mission will change the vehicle to manual Position mode, allowing for quick and easy takeover. The throttle stick should be non-zero when switching to a manual mode to prevent crashes. Manual takeover should be tested in a controlled outdoor environment before attempting a full mission plan. We recommend setting the speed of the drone between waypoints in the mission plan to be 1 m/s, to allow the drone pilot to remain within visual range of the drone without causing high risk to the pilot while traversing uneven ground. Though not required by regulation for non-FPV (first-person view) drones (37), an observer can also help the drone pilot ensure the area is clear before and during each flight.

Permission to fly in the survey area should also be obtained from the land managers, and it must be ensured that the chosen area complies with regulation (see section 4.2).

We also recommend setting up fail-safes in QGroundControl, including automatic landing when the battery level is below a user-defined voltage threshold, when there is a loss of communication with the remote control or telemetry or when a set Geofence (max radius and max altitude of operation space) is breached.

The max altitude and radius of the Geofence should be chosen specifically for the environment being surveyed as the values depend on the altitude that the drone needs to be able to rise to when avoiding obstacles, and the radius that keeps it within line-of-sight and away from restrictions like buildings or people. Regardless of size, having a set Geofence will reduce the likelihood of a flyaway situation and improve safety of operation.

### 4.2 Regulation

UK Civil Aviation Authority (CAA) regulations limit the use of our drone platform to visual line-of-sight (VLOS) operations under an Open Category license. Therefore, for fully autonomous surveying, in which a pilot can set a mission and then allow the drone to complete it without human accompaniment in the field, a Special Category license would be required to allow for beyond visual line-of-sight (BVLOS) operation.

Our drone platform also must be used in accordance with other CAA safety regulations, such as having a Flyer ID for the drone pilot and an Operator ID, keeping a minimum horizontal distance of 50m from people (including those in buildings and transport) that are not involved in the survey, keeping 150m away from residential, recreational, commercial and industrial areas, staying out of Flight Restriction Zones (FRZ) and away from airports, airfields, and other restricted airspaces. Full details are found in the Drone and Model Aircraft Code (38).

## 5. Field deployment and system evaluation

### 5.1 Survey design

We conducted a small-scale survey at the Knepp Estate in West Sussex, to evaluate the ability of our prototype to autonomously avoid real obstacles and to record valid PAM data.

The survey was conducted within a single field (approx. 6.5 acres) in the Southern Block of the Rewilding Project, in order to mitigate the battery life limitations exacerbated by the slower flight speed (1.0 m/s) required to keep the drone within telemetry range, and to ensure the drone pilot can maintain line of sight with the drone. Flight between fields would instead require the drone to fly over trees and, at least temporarily, out of sight of the operator. The survey field was chosen for its accessibility on foot while still having a variety of obstacles to test autonomous obstacle avoidance. Within the field, four locations were chosen as landing sites. The locations chosen had unobstructed landing areas, and maximum separation while still remaining within the same field and within the CAA guidelines. The order of the locations was chosen to maximise the total time the drone was airborne.

We conducted our survey over the course of two days, between the hours of 6am and 4pm. The drone started at location 1 on both days before flying to location 2, recording there for approximately two hours, and so on until it reached location 1 again to record for the final two hours of the survey period (Figure 4). Static AudioMoths, in official AudioMoth cases, were mounted on the side of 1.25 m stakes placed within a few metres of each drone landing site, where they recorded throughout the survey period on both days. The drone AudioMoth was also configured to record throughout the survey period. For all the AudioMoths, we used a sample rate of 48 kHz and a medium gain, with the sleep/record cyclic recording disabled, LED enabled, battery level indication enabled, and the filter disabled.

**Figure 4:**
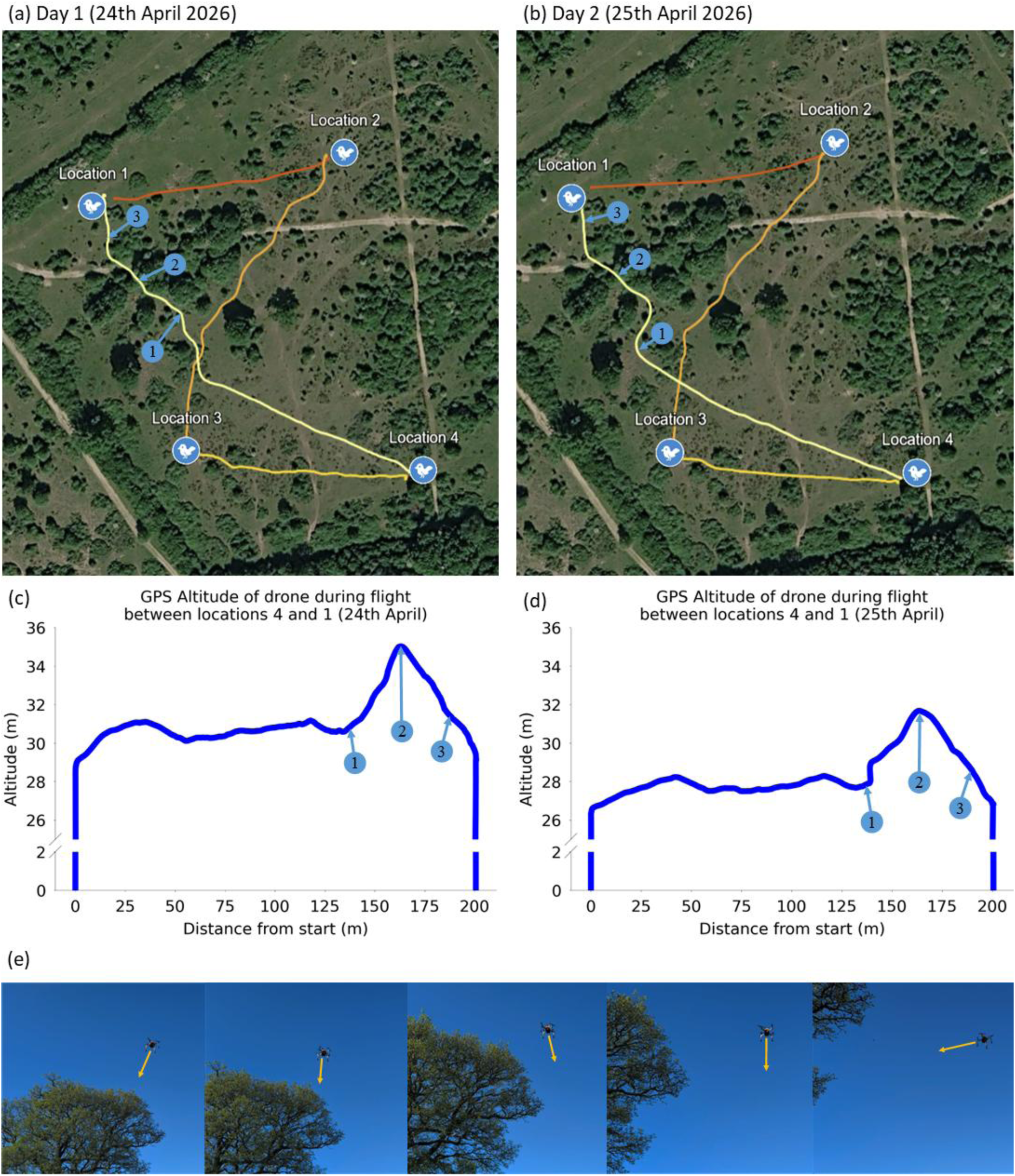
Autonomous flight paths between the four locations of the survey on. (a) Day 1 (24^th^ April) and (b) Day 2 (25^th^ April) and the corresponding altitude maps for (c) Day 1 (24^th^ April) and (d) Day 2 (25^th^ April) with blue numbered markers highlighting the altitude at different points along the flight path. There is clear evidence of obstacle avoidance in three dimensions, but the exact paths taken to avoid the same obstacles on each day are slightly different. A series of snapshots from a video of the drone exhibiting obstacle avoidance behaviour around a tree is shown in (e).

### 5.2 Results

#### 5.2.1 Flight paths with obstacle avoidance

At each location, the drone pilot set the autonomous mission plan to the next location via QGroundControl, with obstacle avoidance enabled. Figure 4 shows the paths taken by the drone between each location across the two days of the survey. Successful obstacle avoidance in the x-y plane is clear in all the paths (Figures 4a, b), but particularly in the second and fourth paths on both days, where trees and tall brush provided direct obstacles for the drone to avoid, which it did so without issue. The altitude plots (Figure 4c, d) for the fourth path on both days also demonstrates obstacle avoidance behaviour along the z-axis, with the altitude remaining relatively constant until the drone reached an area with tall vegetation, where an increase in altitude was required in order to navigate the dense obstacles. While avoidance was successful on both days, the actual paths taken were slightly different between the two days, since the exact paths taken will have been impacted by varying lighting, initial orientation and position of the drone, and wind conditions. Figure 4e shows a series of snapshots from a video of the drone avoiding a tree in the middle of the second flight path (the full video can be found as Supplementary Media). The drone changes direction as it approaches the tree to avoid the canopy, then corrects its heading once it’s clear of the obstacle.

#### 5.2.2 Bird vocalisations

When analysing the audio obtained from the AudioMoths, we used a minimum confidence of 0.5, a sensitivity of 0.5, and an overlap of 2 seconds, in line with the guidelines presented by Pérez-Granados *et al* (39) for community-level analysis. The minimum and maximum bandpass frequencies were kept at the default values of 0 Hz and 1500 Hz respectively. We specified the longitude and latitude of each location, as well as the week (i.e. 15), with a location filter threshold of 0.3 to filter species by location.

The section of the audio shown in Figure 5a that occurs during the drone flight clearly demonstrates why extracting meaningful vocalisations from audio recorded during drone flights is extremely difficult, as the vocalisations are entirely masked by the noise of the motors. Even if the microphone was mounted further from the motors, the noise still significantly masks any other sounds, as seen in Figure 5b, which shows the audio recorded by the nearby static AudioMoth ∼2m away from the drone landing site.

**Figure 5:**
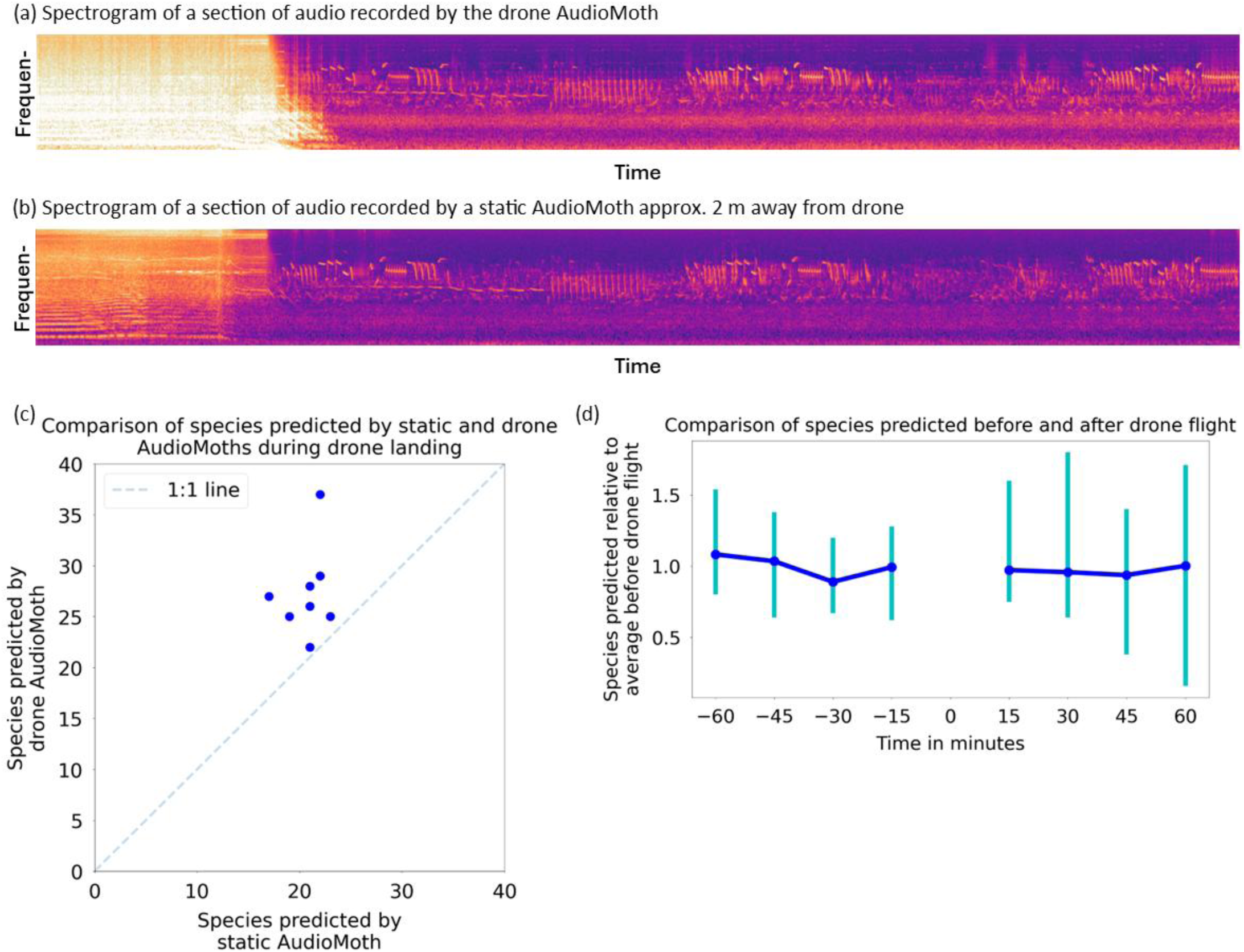
Results of Knepp Estate two-day survey: (a) Spectrogram for section of audio recorded by the drone during landing and after landing, showing the drone noise followed by bird calls. (b) Spectrogram for the same section of audio recorded by the nearby static recorder, showing the drone noise followed by the same bird calls as in (a). (c) Scatter plot comparing species detected by the drone AudioMoth and the static Audiomoths during each drone landing (2 days, 4 locations). The drone Audiomoth detects a greater number of species than the static AudioMoth in every location, as shown by all the points (dark blue) being clearly above the 1:1 line (dotted). (d) Comparison of number of species predicted by the static AudioMoths pre– and post-drone flight, with the points representing the average ratio between the number of species detected in each 15-minute bin and the average number of species detected for the hour before each drone flight. The error bars show the minimum and maximum ratios for each 15-minute bin.

As the way the AudioMoth was mounted on the drone (e.g., its casing, orientation, and nearby components) could impact the number of vocalisations detected, we investigated whether the drone AudioMoth delivered comparable avian biodiversity data to a nearby static AudioMoth. In Figure 5c, we plotted the number of bird species (species richness) predicted by BirdNET for the audio recorded at each location by the drone AudioMoth against the number of bird species predicted by BirdNET for the audio recorded by the corresponding static AudioMoths at each location. All of the points plotted are above the 1:1 line, showing that the number of species detected by the drone AudioMoth exceeds the number of species detected by the static AudioMoth in every location. While this is unexpected, it is likely due to the lack of waterproofing.

The static AudioMoths were secured in standard cases with a waterproof acoustic vent over the microphone rather than an uncovered gap like in our 3D printed case and therefore had greater attenuation than the drone AudioMoth and detected slightly fewer vocalisations. With waterproofing, the number of species detected by the drone AudioMoth would likely become more comparable to the static AudioMoths.

Another factor in determining the efficacy of our approach was the impact of incoming and outgoing drone noise on bird presence at each location. Therefore, we plotted a comparison across all locations where audio was available for 1 hour before and after the drone flight (Figure 5d). We conducted t-tests to determine whether there were statistically significant differences between the distributions of values at 15, 30, 45 and 60 minutes respectively post-drone and the distribution of all values pre-drone. There was no significant difference between the number of species detected 15, 30, 45 and 60 minutes post-drone to the number of species detected pre-drone (p>0.05 in all cases), suggesting that the birds were not significantly disturbed by drone noise.

## 7. Discussion

We successfully demonstrated autonomous operation in an outdoor environment with our modular, open-source platform, suggesting the potential of drone-assisted ecoacoustic surveying using intermittent locomotion in creating far-reaching acoustic surveys into more inaccessible areas. However, there are some aspects of the current design that limit the applicability of the platform in practice. In our trials, a new battery was used for each flight. Reducing power usage (e.g., using more efficient motors or lighter components) could help extend battery life and the reach of the drone, but is unlikely to extend it to a suitable length for long-term surveying. The flight duration of a top-line autonomous commercial drone (DJI Matrice 4) is 49 minutes in a windless environment (40), which would be significantly reduced in real conditions. Our total flight time per field survey was ∼ 8-9 minutes, covering only ∼ 600 m. Therefore, if we assume a roughly linear relationship, the DJI Matrice 4 would be able to cover a theoretical distance of ∼ 4 km, which is likely not sufficient for long-term surveying over large areas and is an overestimate for real, windy conditions. A fully autonomous approach, meanwhile, would be to use wireless charging ground stations at selected sites (41), but over large areas, this would require a number of ground stations to be deployed (42) which would be costly and logistically complex. An alternative autonomous charging method is the addition of solar panels on the drone (43), allowing it to charge while landed during the day. However, the extra weight from the solar panels would increase the overall power consumption of the drone, and this method would preclude usage of the drone in environments where sunlight doesn’t reach the ground, such as forests with dense canopies. In practice, different power solutions are likely necessary for different survey scales and environments. Another potential method of extending the flight range would be to use hybrid drones, which combine fixed-wing and quadcopter technology (44). Fixed-wing drones have a greater flight endurance but require an open space for take-off and landing. Quadcopters can achieve vertical take-off and landing (VTOL) and can therefore operate without large open take-off and landing spaces available.

Combining them can therefore extend flight duration while still maintaining the ability to achieve precise take-off and landing in a range of environments.

Current CAA regulation also limits the utility of our approach without a certified pilot. As per the CAA drone code, the drone must always stay within line-of-sight, which prevents the platform from being used fully autonomously, particularly in inaccessible areas. To conduct beyond-visual-line-of-sight (BVLOS) flight, a designated drone pilot must train to obtain a Level 2 Remote Pilot Certificate (RPC-L2), and the project must apply for a BVLOS operational authorisation, limiting accessibility and cost-efficiency.

Modifications to our drone would also be required for BVLOS flight to enable long-range radio control and telemetry. This can be achieved either using long-range (>40km) radio (45), cellular networks (46), or satellite communication (47).

In our trials, we pre-visited the landing sites to ensure they were suitable. In inaccessible areas, this would not be possible, and therefore the drone would have to autonomously determine whether a proposed landing site is suitable. The PX4 Avoidance software package we use to enable autonomous obstacle avoidance also includes a safe landing planner, allowing the drone to find a safe landing spot by evaluating the ground below to determine if it is flat. We have not tested this capability in our study, but it would be a useful addition towards enabling fully autonomous operation in complex environments where the ground may be significantly uneven or covered with dense vegetation. There has also been significant research into both perching drones and all-terrain landing gear. For example, a study by Lan *et al* (2025) (48) explored the use of tethers, and different perching and disentangling strategies, to enable quadcopters to hang from a branch for long-term monitoring. Similarly, a study by Li *et al* (2025) (49) presented a treecreeper-inspired hardware solution that uses a claw mechanism and supporting “tail” mechanism to allow an aerial robotic platform to grip and perch on tree surfaces. Alternatively, two different studies, by Sarkisov *et al* (2018) (50) and Martynov *et al* (2023) (51) suggest novel rough-terrain landing gear that utilises robotic limbs to provide adaptation to unknown environments and therefore enable safe landing on uneven surfaces.

Depending on the environment being surveyed, either of these approaches (perching vs adaptive landing gear) could be used to allow for robust, long-term monitoring in remote locations.

Our custom drone design has an approximate cost of £1800 for all onboard components. With the addition of spare components, the total approximate cost is £2000, which is comparable to commonly-used off-the-shelf drones with autonomous mission planning capabilities, such as the DJI Mavic 4 Pro (52). The cost could be further reduced by standardising the drone design and mass purchasing components to minimise the per-unit price. Unlike an off-the-shelf drone, however, our custom drone has the flexibility of mounting different sensor modalities, including acoustic sensors. Off-the-shelf designs are typically closed systems that are not easily modified and do not provide any anchor points for external payloads. Moreover, the breakages experienced with our platform have all easily been repaired by us within a few days (usually by replacing inexpensive components such as motors, ESCs, propellers and/or the radio receiver), demonstrating a key benefit of custom drones. Off-the-shelf drones are generally more complex to repair and are usually returned to the manufacturer when damaged, resulting in extra cost and a wait time (for the DJI Matrice series, the average wait time is 3-4 weeks (53)). Using a custom drone also aligns directly with the second principle of Sustainability Robotics (54), an emerging principle that defines minimally invasive sensing methods as a key enabler for sustainable impact with robotic systems. The second principle (universal accessibility) emphasises the need to design robots that extend across “diverse functional, environmental and socio-economic barriers” (54).

Furthermore, our modular design allows for several components to be adapted for different environments and equipment. The landing gear we have designed is appropriate for the environment in which we tested the drone platform, with wide and tall legs that can absorb the impact of landing on hard, uneven ground, and prevent the propellers from getting tangled in low vegetation that is typically found in scrubland during the spring months. However, the landing gear could be swapped out with alternatives depending on the survey environment (e.g., buoyant landing gear to enable landings on water or waterlogged land (55,56) or retractable landing mechanisms to improve shock absorption (57)). The 3D design of the combined case for the onboard computer and acoustic recorder could similarly be adapted for different onboard computer and acoustic recorder models, or other sensor modalities. Even the entire frame could be adapted to be more compact and lighter, for environments where agility is important like dense forests, while keeping the same additional components or replacing them with more compact versions that achieve the same purpose. These design choices all depend on the environment being surveyed and the budget of the project, but modularity allows for researchers to make more fine-grained decisions compared to when using commercial drone models, and to cover a wider range of environments and scenarios.

## 8. Conclusion

In this study, we presented a novel proof-of-concept autonomous drone platform that can use intermittent locomotion to conduct ecoacoustic surveys without degrading audio data with drone noise. We have shown that a modular, open-source approach can enable accurate path planning and obstacle avoidance in real-world outdoor environments and allow for adaptability to new environments and survey designs. In our preliminary trials, we also demonstrated that our approach delivers comparable data to static PAM recording devices. Although regulatory and technical challenges remain, our work paves the way for drones to deliver high quality, fully autonomous, and cost-effective ecoacoustic biodiversity surveys that are accessible and minimally invasive.

### Declaration of generative AI and AI-assisted technologies in the manuscript preparation process

No generative AI or generative AI-assisted technologies were used in the manuscript preparation process.

## Author Note

We have no known conflict of interest to disclose.

## CRediT Author contributions

Milica Ostojic: Conceptualisation, data curation, formal analysis, investigation, methodology, software, visualisation, writing – original draft, writing – review and editing.

Oscar Pang: Investigation, methodology, resources, writing – review and editing.

Matt Phelps: Resources, writing – review and editing.

Mirko Kovac: Conceptualisation, funding acquisition, resources, writing – review and editing.

Sarab Sethi: Conceptualisation, funding acquisition, methodology, supervision, visualisation, writing – review and editing.

## Data Availability Statement

The BirdNET detections for the audio collected in this study are available on Zenodo at https://doi.org/10.5281/zenodo.22920446. The raw audio is unavailable due to privacy reasons.

## Supporting information

Supplemental Media

## Acknowledgements

We thank the Knepp Estate Rewilding Project for granting us permission to conduct fieldwork on site, and for their support with the project.

