## Supplemental Table for "Fly, land, listen: Autonomous intermittent locomotion enables scalable low-noise drone ecoacoustic surveys"

Supplementary Table

| Model | Link to purchase |
|---|---|
| (3DXR) HexSoon – EDU-450 | https://www.3dxr.co.uk/multirotor-c3/multirotor-frames-c97/hexsoon-edu-450-v2-p5176 |
| (3DXR) T-Motor T1045 Self-locking Props | https://www.3dxr.co.uk/multirotor-c3/multirotor-props-c265/t-motor-t1045-self-locking-props-pair-p3209 |
| (Holybro) Brushless Motor 2216-920KV | https://www.flyingtech.co.uk/product/holybro-brushless-motor-2216-920kv-cw-ccw/ |
| 3.5 mm bullet connectors | https://amzn.eu/d/0hlYFR9t |
| (3DXR) T-Motor – Air 20A (V2) ESC 3-4S | https://www.3dxr.co.uk/fixed-wing-c27/fixed-wing-escs-c52/t-motor-air-20a-v2-esc-3-4s-no-bec-p2791 |
| (Holybro) Pixhawk 6C | https://holybro.com/products/pixhawk-6c |
| (Holybro) PM06 Power module | Purchased with Pixhawk 6C, or separately: https://holybro.com/products/micro-power-module-pm06-v2 |
| (Holybro) Cable set including PWM board, pin cables and USB cable | Purchased with Pixhawk 6C, or separately: https://holybro.com/products/pixhawk-6c-cable-set |
| (Holybro) M9N GPS (Standard) | https://holybro.com/products/m9n-gps |
| (Holybro) SiK Telemetry Radio V3 (100mW 433Mhz) | https://holybro.com/products/sik-telemetry-radio-v3 |
| (Discontinued) FrSky Taranis X-Lite Pro 2.4GHz ACCESS Radio Transmitter | N/A (any radio controller compatible with the ACCESS protocol should be sufficient) |
| (FlyingTech) FrSky R9M Lite Pro Long Range Transmitter Module – ACCESS | https://www.flyingtech.co.uk/product/frsky-r9m-lite-pro-long-range-transmitter-module-access/ |
| (FlyingTech) FrSky R9 Slim + OTA 868MHz 16CH SBUS Receiver with ACCESS Protocol | https://www.flyingtech.co.uk/product/frsky-r9-slim-ota-868mhz-16ch-sbus-receiver-with-access-protocol/ |
| (3DXR) Benewake – TFmini-S Micro LiDAR – (12m range) – UART | https://www.3dxr.co.uk/sensors-c5/lidar-range-and-flow-sensors-c4/benewake-tfmini-s-micro-lidar-12m-range-p5142 |
| (HobbyRC UK) GNB 3000mAh 4S 100C LiPo Battery | https://www.hobbyrc.co.uk/gnb-3000mah-4s-100c-lipo-battery |
| (3DXR) Nylon strap with plastic buckle | https://www.3dxr.co.uk/building-c23/tapes-c130/velcro-c350/3dxr-nylon-strap-with-plastic-buckle-2-pcs-p5217 |
| Intel RealSense D455 | https://store.realsenseai.com/buy-intel-realsense-depth-camera-d455.html |
| Cable Matters 2-Pack, USB to USB C Coiled Cable | https://amzn.eu/d/0gnjH5Jl |
| (Discontinued) NVIDIA Jetson Orin Nano Developer Kit | New model: https://www.siliconhighwaydirect.com/product-p/945-13766-0005-000.htm |
| (Discontinued) Crucial P3 1TB M.2 PCIe Gen3 NVMe Internal SSD / (Alternative) Crucial P310 1TB SSD M.2 2280 NVMe PCIe Gen4 | https://amzn.eu/d/0dxNA182 |
| FTDI Serial Breakout Module Programmer – 5V/3.3V (USB to Serial TTL) | https://www.flyingtech.co.uk/product/ftdi-serial-breakout-module-programmer-5v-3-3v-usb-to-serial-ttl/ |
| AudioMoth | https://www.openacousticdevices.info/product-page/audiomoth-1-2-0 |
